# Additive genes underlie Mendelian inheritance and contribute to heterosis

**DOI:** 10.64898/2026.11.12.766085

**Authors:** Zhiwu Dan, Yunping Chen

**Author notes:** Corresponding author: Zhiwu Dan, National Technology Innovation Center for Saline-Alkali Tolerant Rice, Sanya 572024, China. These authors contributed equally.

## Abstract

Mendel proposed the laws of segregation and independent combination in his study. However, he focused on “dominant” characters but left “intermediate” ones. Using genotypic and phenotypic data for four representative genes, we reveal connections between the “intermediate” and “dominant” characters. We demonstrate that the fundamental inheritance patterns of these genes are additive effects on both component and compound traits. Additive effects involving at least two genes generate non-additive effects in hybrids. For compound traits such as grain yield, their inheritance depends on differential patterns of component traits in parents; only parents with optimal balances of reciprocal differences in component traits produce overdominant effects, which is important for hybrid breeding.

## Introduction

In 1865, Gregor Mendel spoken in a lecture that “*With some of the more striking characters, those, for instance, which relate to the form and size of the leaves, the pubescence of the several parts, etc*., *the* ***intermediate***, *indeed*, ***is nearly always to be seen***; *in other cases, however, one of the two parental characters is so preponderant that it is difficult, or quite impossible, to detect the other in the hybrid*.” (http://www.mendelweb.org/homepage.html) (Mendel, 1866). He did not focus on the “***more striking characters***” but on the “***other cases***”, among which one parental character is “***so preponderant****”* that the other is difficult or impossible to detect in F_1_ hybrids. He used the terms “dominant” and “recessive” and the letters “*A*” and “*a*” to describe corresponding factors, respectively. He found that the investigated phenotypes of F_1_ hybrids with genotype “*Aa*” were identical to those of the parental “*AA*” and showed that the ratio of dominant to recessive characters was 3 to 1 in the F_2_ generation, which is the law of segregation in Mendelian inheritance. He further proposed that each pair of differential characters is independent of other ones in egg and pollen cells, which is the law of independent combination.

Compared with the “***other cases***”, the “***more striking characters***” are more common characters that were largely uninvestigated in Mendel’s research. However, Mendel used “intermediate” to describe these characters, referring to an additive effect represented by the average level of the parents. The genetic underlying of these intermediate characters remain unsolved, and the connections between the intermediate and dominant characters are unknown. In this study, we use genotypic data of four representative genes and corresponding phenotypic data to investigate these details.

## Results and discussions

Rice yield is a compound trait and is mainly determined by four component traits: grain number per panicle (GNP), thousand-grain weight (TGW), tiller number per plant (TPP), and seed setting rate (SSR). These component traits are themselves composed of sub-traits, such as primary branch number (PBN) and grain length (GL). In addition, rice yield is associated with other yield-related traits, including plant height (PH) and heading date (HD).

*OsMADS1* is a gene that mainly regulates GL, TGW, and grain quality (Wang et al., 2024). Its mutants (designated *A*^+^*A*^+^ in this study) exhibit larger GL and TGW and delayed HD, but show no significant differences from wild-type plants (designated *A*^-^ *A*^-^ in this study) in GNP, SSR, TPP, panicle length (PL), and PH (Figure 1a). When this gene is in the heterozygous state (designated *A*^+^*A*^-^) in BC_2_F_2_ lines, it shows additive effects on GL, TGW, and YPP (Figure 1b). Similarly, heterozygous *OsMADS1* is intermediate between the parents for GL and TGW in F_6_ RILs and for GL, TGW, and YPP in BC_4_F_2_ lines (Liu et al., 2018; Wang et al., 2019).

**Figure 1.**
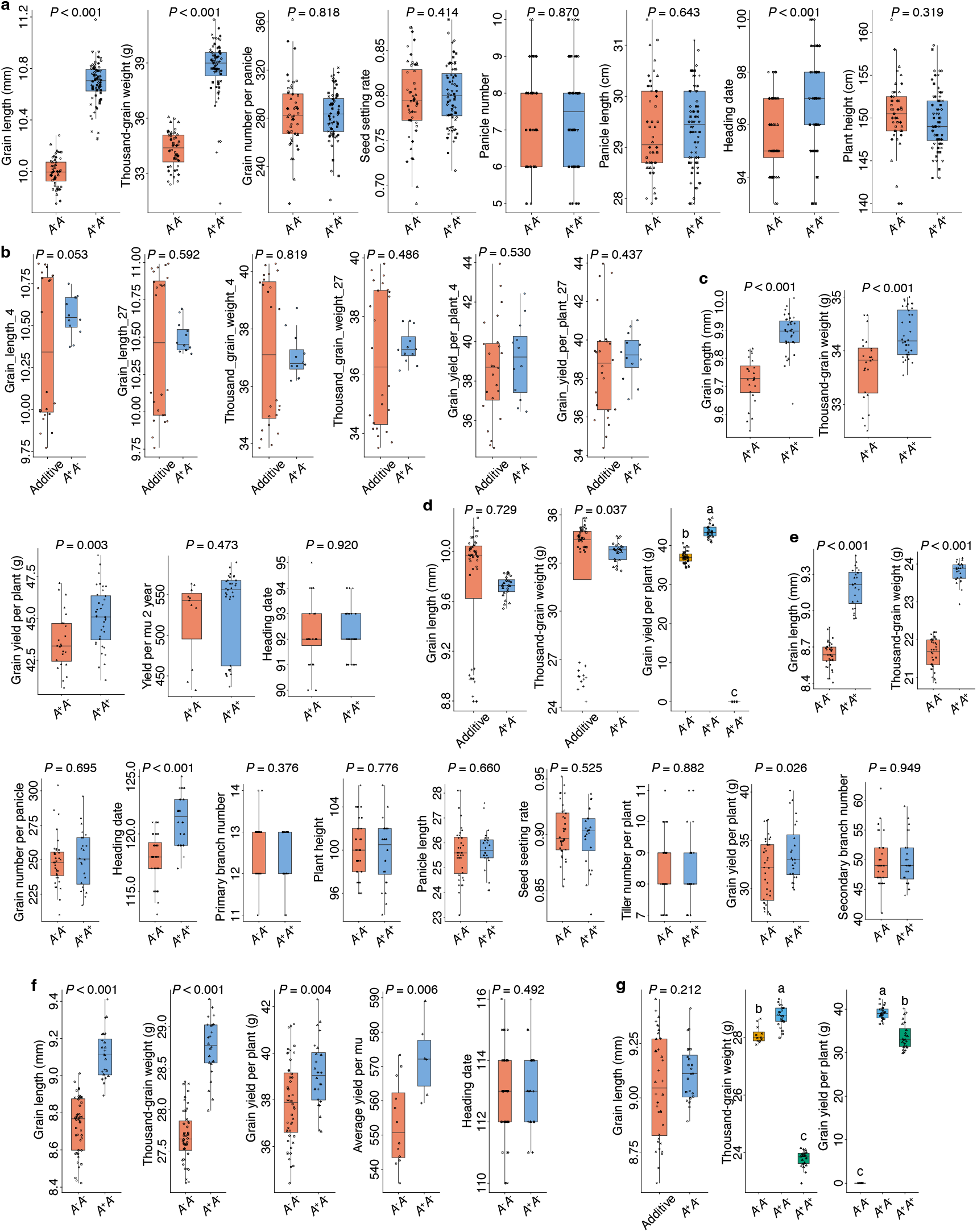
Comparisons of homozygous and heterozygous *OsMADS1*. (a), Trait values for different homozygous genotypes of *OsMADS1* in Fuhui676. (b) Trait values of heterozygous *OsMADS1* and corresponding additive effects in BC_2_F_2_ lines. (c) Trait values of wild-type and *OsMADS1*-mutant GLY676. (d) Inheritance patterns of *OsMADS1* for component and compound traits of GLY676. (e) Trait values for different homozygous genotypes of *OsMADS1* in R534. (f) Trait values of wild-type and *OsMADS1*-mutant QYSM. (g) Inheritance patterns of *OsMADS1* for component and compound traits of QYSM.

In F_1_ hybrids derived from crosses between restorer and sterile lines, the mutant F_1_ hybrids with homozygous *OsMADS1* (*A*^+^*A*^+^) have larger GL, TGW, and YPP and comparable HD compared to heterozygous ones (*A*^+^*A*^-^) (Figure 1c). When F_1_ hybrids heterozygous for *OsMADS1* were compared to their corresponding parents, an additive effect is observed for GL (Figure 1d). However, a partially dominant effect—between the additive effect and the better parent—is found for TGW. An overdominant effect— surpassing the better-parent—is found for YPP. In another F_1_ hybrid with the restorer line carrying *OsMADS1* mutations, consistent with results in Figure 1a, homozygous mutant lines (*A*^+^*A*^+^) display larger GL, TGW, and YPP and elongated HD, but no significant differences in GNP, PBN, PH, PL, SSR, TPP, or secondary branch number compared with wild-type lines (*A*^-^*A*^-^) (Figure 1e). In F_1_ hybrids, those heterozygous for *OsMADS1* (*A*^+^*A*^-^) have larger GL, TGW, and yield and comparable HD compared with wild-type hybrids (*A*^-^*A*^-^) (Figure 1f). Notably, when compared with the homozygous parents, an additive effect is observed for GL, whereas overdominant effects are observed for both TGW and YPP (Figure 1g), indicating a shift from additive to overdominant effects from component to compound traits.

*IPA1* is a gene regulating multiple agronomic traits, including TPP, GNP, YPP, and PH (Song et al., 2022). Lines heterozygous for a single *IPA1* mutation display additive effects for TPP, GNP, YPP, PH, TGW, SSR, panicle weight, root diameter, and stem diameter (Figure 2a). Consistent with the findings for both *OsMADS1* and *IPA1*, additive effects are found for PH and amylose content in lines heterozygous for *Ghd7* (Figure 2b), which regulates PH, YPP, and HD (Lou et al., 2026). In foxtail millet, heterozygous lines for the green-revolution gene *SiSD1*, a gibberellin biosynthesis gene that regulates PH and TPP (Wang et al., 2026), show additive effects for GA1, GA3, PH, and TPP (Figure 2c). However, an overdominant effect is found for YPP in F_1_ hybrids heterozygous for *SiSD1*, confirming the shift in inheritance patterns from component to compound traits across crops.

**Figure 2.**
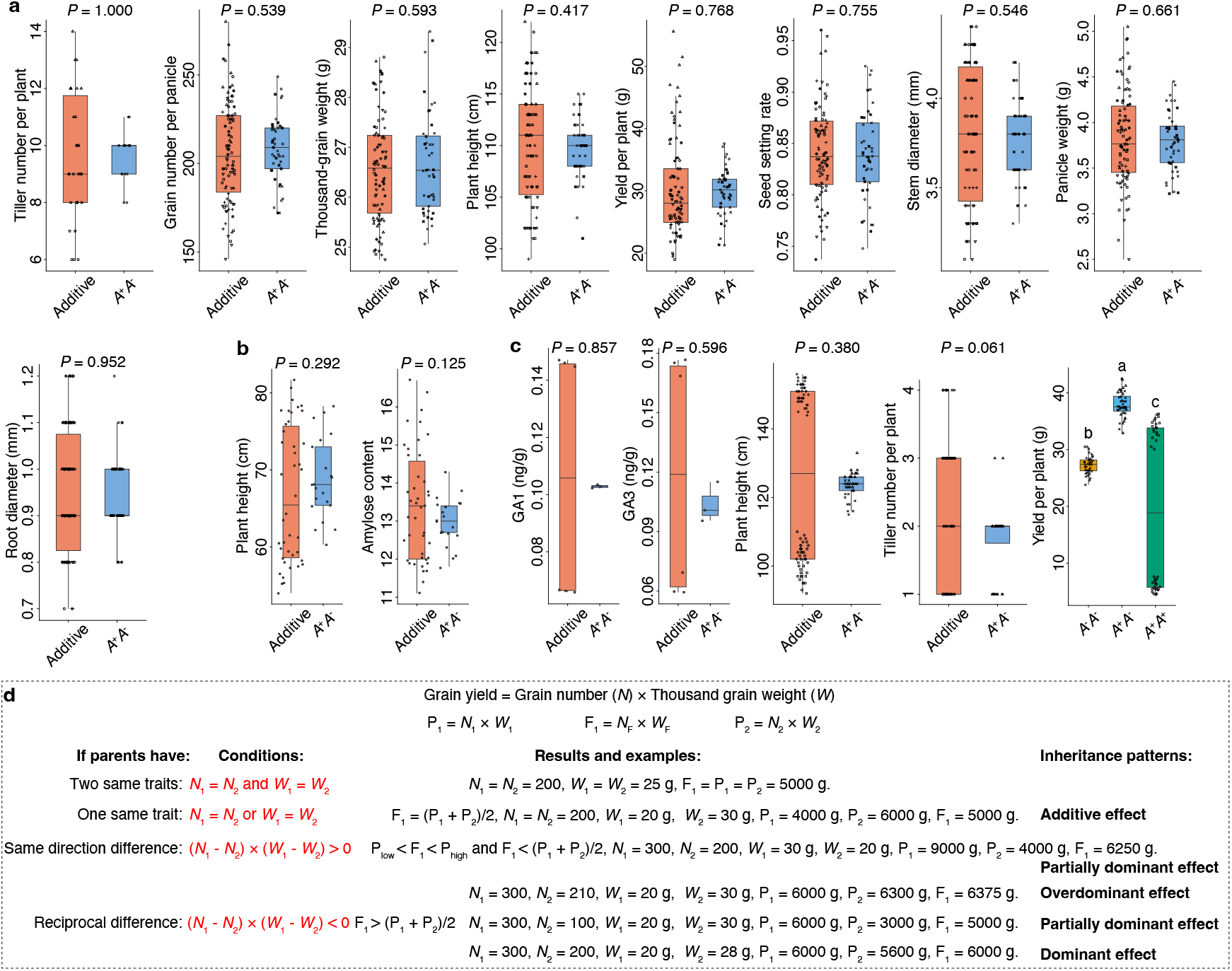
Comparisons of three genes in homozygous or heterozygous states and the contributions of additive genes to component and compound traits. (a) Trait values of heterozygous *IPA1* and corresponding additive effects. (b) Trait values of heterozygous *Ghd7* and corresponding additive effects. (c) Gibberellin content and trait values of heterozygous *SiSD1* and corresponding additive effects. (d) Rationale for additive effects to component and compound traits.

Based on these results and our previous findings (Dan et al., 2026a; Dan et al., 2026b), we conclude that the additive effect is the fundamental inheritance pattern and that other non-additive effects are derived from it. We propose a simplified rationale in Figure 2d. To facilitate understanding, grain yield is assumed to consist of only two compound traits: grain number (*N*) and thousand-grain weight (*W*). The grain yield of two parents and F_1_ hybrids are given by the equations P_1_ = *N*_1_ × *W*_1_, P_2_ = *N*_2_ × *W*_2_, and F_1_ = *N*_F_ × *W*_F_, respectively. We classify the differences in *N* and *W* between the two parents into four types: two same traits, one same trait, same-direction difference, and reciprocal difference. Under the additive-effect rule, the first three types yield expected outcomes, including additive and partially dominant effects. The fourth type encompasses three inheritance patterns: partially dominant, dominant, and overdominant effects. Notably, if the two parents have the same yield but also show reciprocal differences in *N* and *W*, an overdominant effect, namely positive heterosis, will undoubtedly be observed for yield in F_1_ hybrids. Thus, additive effects of at least two genes produce non-additive inheritance patterns in F_1_ hybrids.

Mendel used the words “dominant” and “recessive” to describe characters. In fact, those “dominant” characters are likely the integrated results of at least two additive genes. Therefore, the representing of these genes as “*A*” and “*a*” may be in appropriate; using “*A*^+^” and “*A*^-^”, referred to as *A* positive and *A* negative, respective, is more accurate.

Because F_1_ hybrids carrying a single heterozygous gene, or backcross/RILs with heterozygous genotypes at the target genes, exhibit additive effects for both component and compound traits, the inheritance patterns of compound traits in hybrid crops depend on the differential patterns of component traits in their parents. Only the parents that have reciprocal differences in component traits can produce overdominant effects for compound traits. Optimal balances in the reciprocal differences of component traits ensure the realization of heterosis for compound traits such as grain yield.

## Materials and methods

No transgenic lines were generated, and no phenotypic data were recorded in this study. All genotypic and phenotypic data were obtained from previous studies and used for statistical analyses (Wang et al., 2019; Song et al., 2022; Wang et al., 2024; Lou et al., 2026; Wang et al., 2026). Additive effects were represented by the averages of parental values. Figures were prepared using grafify online (Shenoy, 2026). Independent-samples *t*-tests, Welch’s *t*-tests, and analysis of variance (ANOVA) were used for comparative analyses.

## Author contributions

Z.W.D. designed the study, collected and analyzed data, prepared figures, and wrote the manuscript. Y.P.C. analyzed data, prepared figures, and wrote the manuscript.

## Acknowledgement

This work was supported by 2026 National Innovation Center for Salt-Tolerant and Alkali-Tolerant Rice Project (ZDYF2026GCZX001). The authors are grateful to Dr. Tianshun Zhou for helpful discussions.

